# SARS-CoV-2 Spike Protein Reduces Burst Activities in Neurons Measured by Micro-Electrode Arrays

**DOI:** 10.1101/2023.04.24.538161

**Authors:** Melanie Salvador, Noah Tseng, Camdon Park, Grace Williams, Arianne Vethan, Grant Thomas, John Baker, Joseph Hemry, Emma Hammond, Paige Freeburg, Guan-Wen Chou, Nick Taylor, Yi-Fan Lu

**Affiliations:** Biology Department, Westmont College, CA, 93108; Department of Computer Science, North Carolina State University, NC 27695

## Abstract

SARS-CoV-2 caused a large-scale global pandemic between 2020 and 2022. Despite efforts to understand its biology and mechanisms of pathogenicity, the viral impact on the neurological systems remains unclear. The main goal of this study was to quantify the neurological phenotypes induced by SARS-CoV-2 spike protein in neurons, measured by *in-vitro* multi-well micro-electrode arrays (MEAs). We extracted the whole-brain neurons from the newborn P1 mice and plated them on multi-well micro-electrode arrays and administered purified recombinant spike proteins (S1 and S2 subunits respectively) from the SARS-CoV-2 virus. The signals from the MEAs were transmitted from an amplifier to a high-performance computer for recording and analysis. We used an in-house developed algorithm to quantify neuronal phenotypes. Among all the phenotypic features analyzed, we discovered that the S1 protein of SARS-CoV-2 decreased the mean burst numbers observed on each electrode; This effect was not observed for the spike 2 protein (S2) and could be rescued by an anti-S1 antibody. Finally, our data strongly suggest that the receptor binding domain (RBD) of S1 is responsible for the reduction of burst activities in neurons. Overall, our results strongly indicate that spike proteins may play an important role in altering neuronal phenotypes, specifically the burst patterns, when neurons are exposed during early development.

## Introduction

SARS-CoV-2 is the pathogenic agent responsible for the coronavirus disease 19 (COVID-19) pandemic. SARS-CoV-2 shares many common features with SARS-CoV that caused a brief global pandemic in 2002, which caused belief in their seemingly common ancestry (Lan et al., 2020). In the recent pandemic, prevalent neurological symptoms observed in hospitalized COVID-19 patients included ataxia, impaired consciousness, neuralgia, and further musculoskeletal symptoms (Kambiz et al., 2020). The SARS-CoV-2 structure primarily consists of membrane (M), envelope (E), spike (S), and nucleocapsid (N) proteins (Mohamadian et al., 2021). The focus of this study addresses the effects of the S protein. The S protein assists in the attachment and entry of the virus into the host cell (Takeda, 2022). Two subunits make up the protein, each with its own function (Cui et al., 2022). The S1 subunit is responsible for receptor binding, and the S2 subunit facilitates membrane fusion (Demers-Mathieu et al., 2021). The protein is structured as a trimer (three receptor binding domains), with the SARS-CoV-2 strain enabling the trimeric structure to open a pathway via pneumocytes or host cells (Jessie et al., 2021). The S proteins of SARS-CoV-2 and SARS-CoV share a sequence identity of 77%, with the predominant difference being solely in the overall 3D structure and its specific binding affinity of the SARS-CoV-2 S protein to the ACE2 on the host receptor (Kambiz et al., 2020).

The S1 subunit binds to the ACE2 receptor which is found on the surface of intestinal and lung cells (Chuang et al., 2022). Within the brain, ACE2 can be seen to be expressed predominantly in the brain stem and regions focusing specifically on regulating cardiovascular function and blood pressure. (Santos et al., 2018). The proposed mechanism for viral attachment involves several steps that take advantage of two cleaving sites on the protein (Wang et al., 2020). One is located between the S1 and S2 subunit, and the other is on the S2 subunit (Jaimes et al., 2020). In the first step a proprotein convertase, predominantly furin, cleaves at the S1/S2 cleavage site (Wrobel et al., 2020). It may be noted that this cleavage is specific to SARS-CoV-2 and does not occur in the original SARS virus, likely contributing to the higher infectious properties of SARS-CoV-2 relative to the original SARS strain (Takeda, 2022). This cleavage destabilizes the S protein, increasing its affinity to bind to ACE2 (Benton et al., 2020; Walls et al., 2020). A D614G mutation allows rotation at that location to adopt the open conformation so that the binding site is exposed (Yurkovetskiy et al., 2020). Finally, cleavage at the S2 binding site performed by transmembrane protease serine 2 (TMPRSS2) results in membrane fusion (Hoffmann et al., 2020). Here, three previously buried hydrophobic fusion peptides in S2 become exposed and inserted into the target host membrane, inducing membrane fusion (Jackson et al., 2020).

To further understand the mechanism of SARS-CoV-2 on the neural system, we utilized multi-well microelectrode arrays (MEAs) to assess the neurological phenotypes *in-vitro*. MEAs contain arrays of electrodes that detect local field potentials that are generated by electrical signals from electrically active cell types, such as neurons or cardiomyocytes (Lu et al, 2015; Cao et al., 2012). In this study, multi-well micro-electrode (MEA) arrays that contained 24 wells and a 4-by-4 array (16 electrodes) per well were used to assess the impacts of SARS-CoV-2 spike proteins on neurons. Utilization of the MEA allows for the assessment of neuronal activity through the detection of changes in extracellular field potentials, which reflect the intrinsic action potentials of cultured neurons. The MEA possesses the ability to analyze electric signals of a population of neurons for an extended period of time (120,000 cells in this experiment for two weeks), exceeding the capability of a patch-clamp experiment which only observes individual cells. The prolonged measurement over a population of neurons is an excellent source to determine the group behavior without constant perturbation, and is somehow analogous, although not identical, to that of the activities of the brain (Hales et al., 2010). MEAs have been extensively used to assess the neurotoxicity of compounds on the behavior of neurons (Valdivia et al., 2014; Wallace et al. 2015; Robinette et al., 2011).

In this study, we utilized MEA as an objective and non-invasive measure to model whether spike proteins induce neurological phenotypes under various conditions. This study may put forth a better understanding of the molecular mechanism of how the SARS-CoV-2 virion influences the neuronal system. Our study shows that the S1 protein reduces the formation of robust bursts, and it is only potent when the neurons are exposed early during the developmental stage. We further demonstrate that the reception-binding domain (RBD) of S1 alone is sufficient to reduce burst activities in the neuronal population. Our study offers new insight into the biology of SARS-CoV-2 and provides an encouraging platform to assess neurological damages caused by SARS-CoV-2 proteins.

## Materials and Methods

### MEA Plate Preparation

Sterilization of a 24-well MEA plate (AutoMate Scientific) was carried out by submerging the plate in 70% ethanol for 5 minutes. The plate was then air-dried for three hours or until all the remaining ethanol was completely evaporated. The coating process was performed by applying poly-D-lysine (50 μL/mL) and the plate was incubated at 37 degrees for one hour. The plate was then washed with sterile water and air-dried before cell plating.

### Disassociation and culturing of primary mouse cortical neurons

Primary mouse cortical neurons were obtained from the newborn P1 mice. 24-well MEA plates were coated prior to the seeding of disassociated neurons. Dissection and disassociation protocols were adopted and optimized from the published work of cortical culture on MEA. The cerebral cortex was removed in Hank’s Balanced Salt Solution (HBSS) buffer at ice--cold temperatures, and was thereafter mechanically disassociated by scissors to 1 to 2 mm in size. The tissue was subsequently subjected to a papain and DNase mixture for 30 minutes in a 37 degrees Celsius water bath. Cells were centrifuged at 200 RCF for 5 minutes and thereafter titrated by a P1000 pipette tip. 120,000 cells were seeded on top of each well covered by electrodes. Plating was done in a randomized pattern to control for confounding between spatial effects on the plates.

### Cell Culture and Spike Protein Treatment

Neurobasal--Plus media (Thermo Scientific) supplied with B-27, 10 mM HEPES, 100 U/mL Penicillin Streptomycin, and 2 mM Glutamax were used to support the development of the neuronal culture. The media was partially changed every other day to maintain the health of the culture. Recombinant SARS-CoV-2 Spike Protein, S1, and S2 subunits respectively, were obtained from RayBiotech and stored at −80 degrees Celsius. On the day of neuronal cell isolation, the spike proteins were diluted and added to the culture media at the desired concentrations. To neutralize the S1 protein, 50 ng of human anti-S1 antibody (ACROBiosystems) was mixed with S1 protein and incubated at room temperature for three hours with gentle agitation before adding it to the culture media.

### MEA plate recording and Spike Quantification

Spikes were detected on a 24-well MEA plate on the MED64 Presto System (AutoMate Scientific) and the MEA Symphony Software. The threshold for calling a spike from the waveform was 500% of the standard deviation generated from the background noise. Time stamps were recorded for the spikes that surpassed the threshold. Spikelist.cvs files with the timestamp information were exported and further analyzed in the RStudio (R-project.org). The maximum resolution of the time stamps was 0.01 millisecond (ms) which was able to distinguish two different spikes.

### Burst Definition

Bursts were defined by an in-house developed algorithm in RStudio. The threshold to define a burst was set at contentious inter-spike intervals of less than 80 ms. Qualified bursts were recorded in a text file and exported for further analysis. After a qualifying burst was determined by the algorithm, parameters such as the timestamp, duration, spike numbers, and spike frequencies in a burst were calculated and recorded. Heatmaps of the burst data were plotted in GraphPad Prism (Insightful Science, Inc), and the density estimation and distribution graphs were generated using the density and plot functions in RStudio (R-project.org).

### Statistical tests

Mean, Standard Error, and 95% confidence interval (CI) were calculated in RStudio (RStudio.org). GraphPad Prism (Insightful Science, Inc) was used to perform statistical tests. Unpaired two-tailed t-tests were conducted with a significance level set at 0.05.

## Results

### Purified S1 protein reduces burst activities in neurons

Although neurological symptoms have been reported in some, not all, COVID patients, the exact mechanism of how viruses affect neuronal cells remains unclear, and thus an area of exploration. Here, we hypothesized that the spike protein of SARS-CoV-2 is responsible for the neuronal phenotypes without the need for infection by the whole infectious virion. In this study, two subunit parts of the spike protein, the S1 subunit, and the S2 subunit, were tested separately to assess whether they induced any neurological phenotypes measured by the MEAs. We obtained recombinant SARS-CoV-2 spike protein, S1 subunit, and S2 subunits, respectively, and treated neurons isolated from the P1 mice on day 0 (day 0 was defined as the day of cell preparation) (Figure 1). Spike protein concentrations were maintained at the same level in the culture media and data from multi-electrode arrays were collected over a two-week period. We analyzed the data through an in-house developed algorithm in RStudio. Specifically, we quantified features such as the number of bursts per electrode, duration, frequency, and spike number per burst based on the treatment condition. Among all the features we investigated, we identified the number of bursts per electrode as the most prominent feature that distinguished the spike protein-treated wells and the control. The S1 subunit significantly reduced the number of bursts per electrode (Table 1 and Figure 2A-C), although this type of reduction was not as noteable in the S2 subunit (Figure 2D-F). Our result suggests that S1 is responsible for reducing burst activities of populations of neurons if cells are exposed early (day 0) during the developmental course.

**Figure 1.**
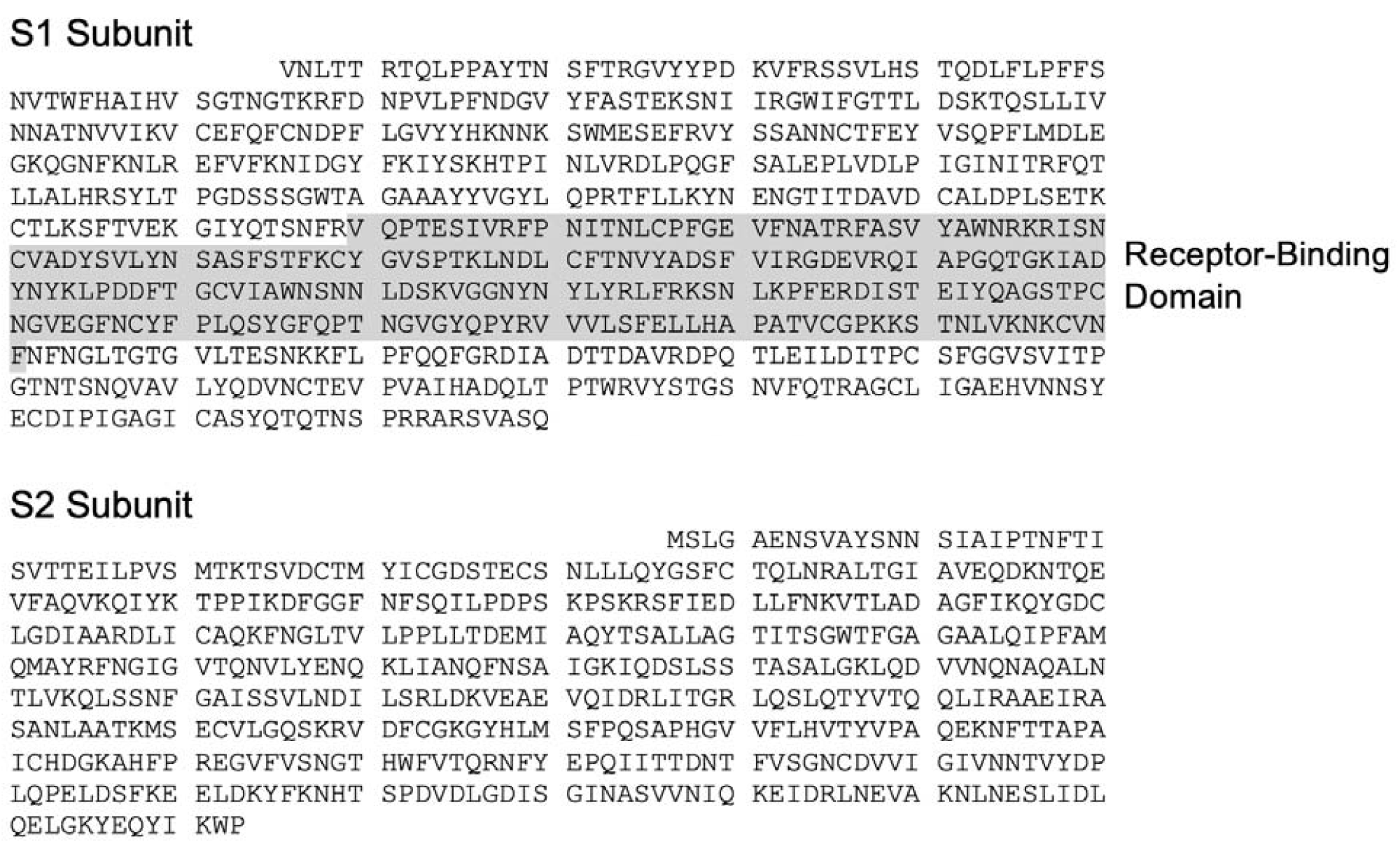
Amino acid sequences of SARS-CoV-2 Spike 1 and 2 used in this study. NCBI accession number QHD43416.

**Table 1.**
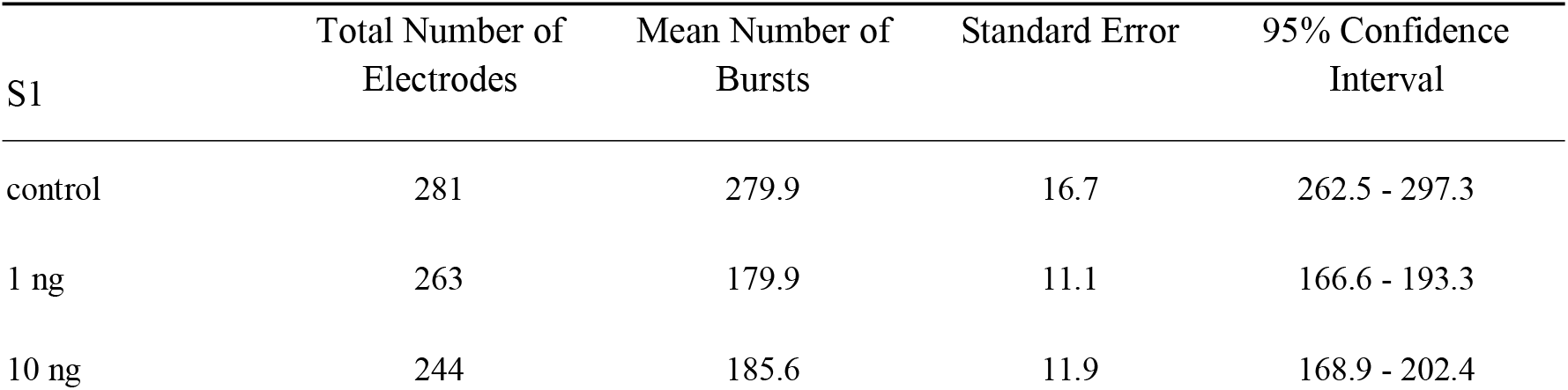
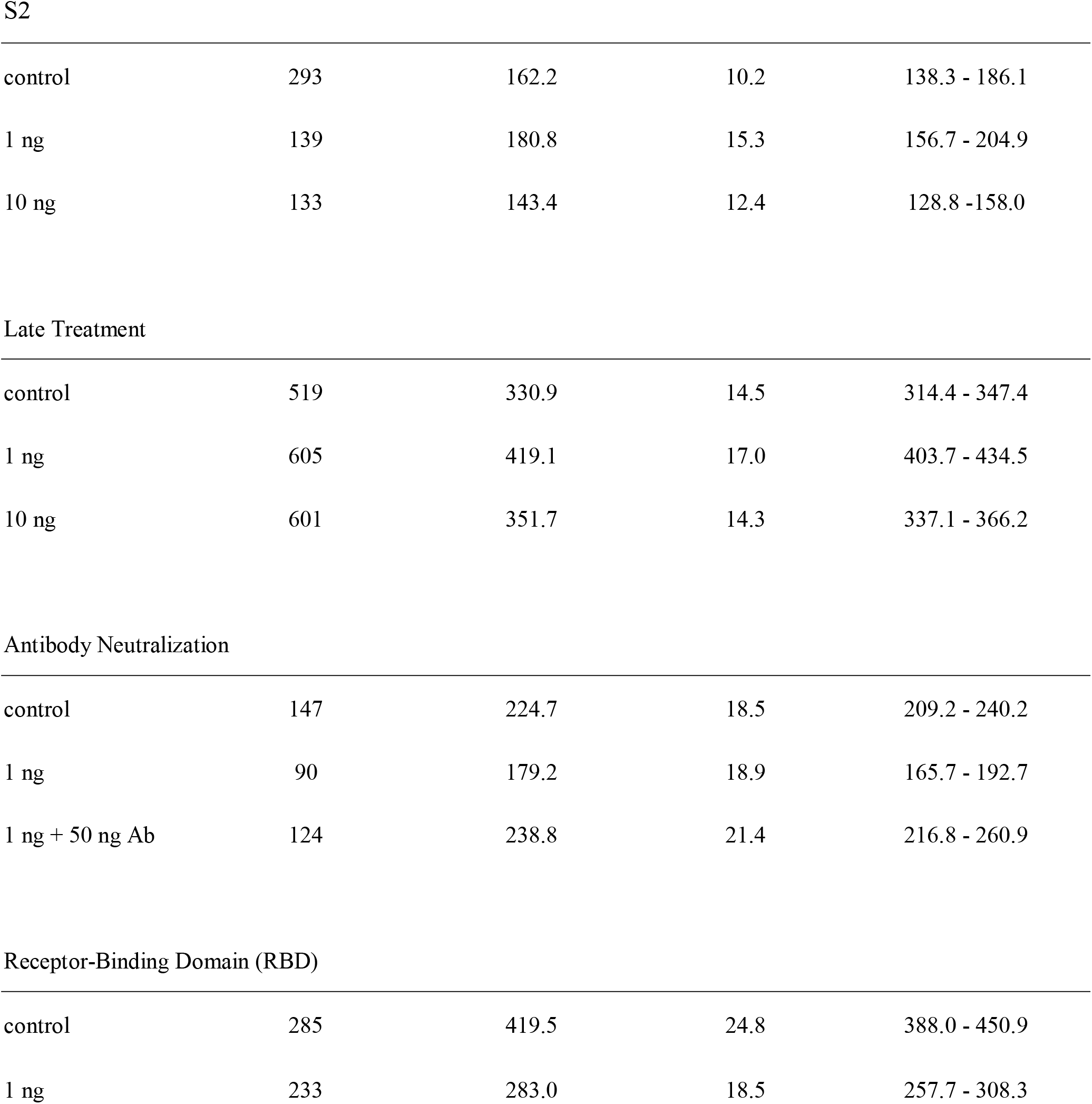
Summary Statistics in This Study.

| S1 | Total Number of Electrodes | Mean Number of Bursts | Standard Error | 95% Confidence Interval |
| --- | --- | --- | --- | --- |
| control | 281 | 279.9 | 16.7 | 262.5 - 297.3 |
| 1 ng | 263 | 179.9 | 11.1 | 166.6 - 193.3 |
| 10 ng | 244 | 185.6 | 11.9 | 168.9 - 202.4 |
| S2 |  |  |  |  |
| control | 293 | 162.2 | 10.2 | 138.3 - 186.1 |
| 1 ng | 139 | 180.8 | 15.3 | 156.7 - 204.9 |
| 10 ng | 133 | 143.4 | 12.4 | 128.8 - 158.0 |
| Late Treatment |  |  |  |  |
| control | 519 | 330.9 | 14.5 | 314.4 - 347.4 |
| 1 ng | 605 | 419.1 | 17.0 | 403.7 - 434.5 |
| 10 ng | 601 | 351.7 | 14.3 | 337.1 - 366.2 |
| Antibody Neutralization |  |  |  |  |
| control | 147 | 224.7 | 18.5 | 209.2 - 240.2 |
| 1 ng | 90 | 179.2 | 18.9 | 165.7 - 192.7 |
| 1 ng + 50 ng Ab | 124 | 238.8 | 21.4 | 216.8 - 260.9 |
| Receptor-Binding Domain (RBD) |  |  |  |  |
| control | 285 | 419.5 | 24.8 | 388.0 - 450.9 |
| 1 ng | 233 | 283.0 | 18.5 | 257.7 - 308.3 |

**Figure 2.**
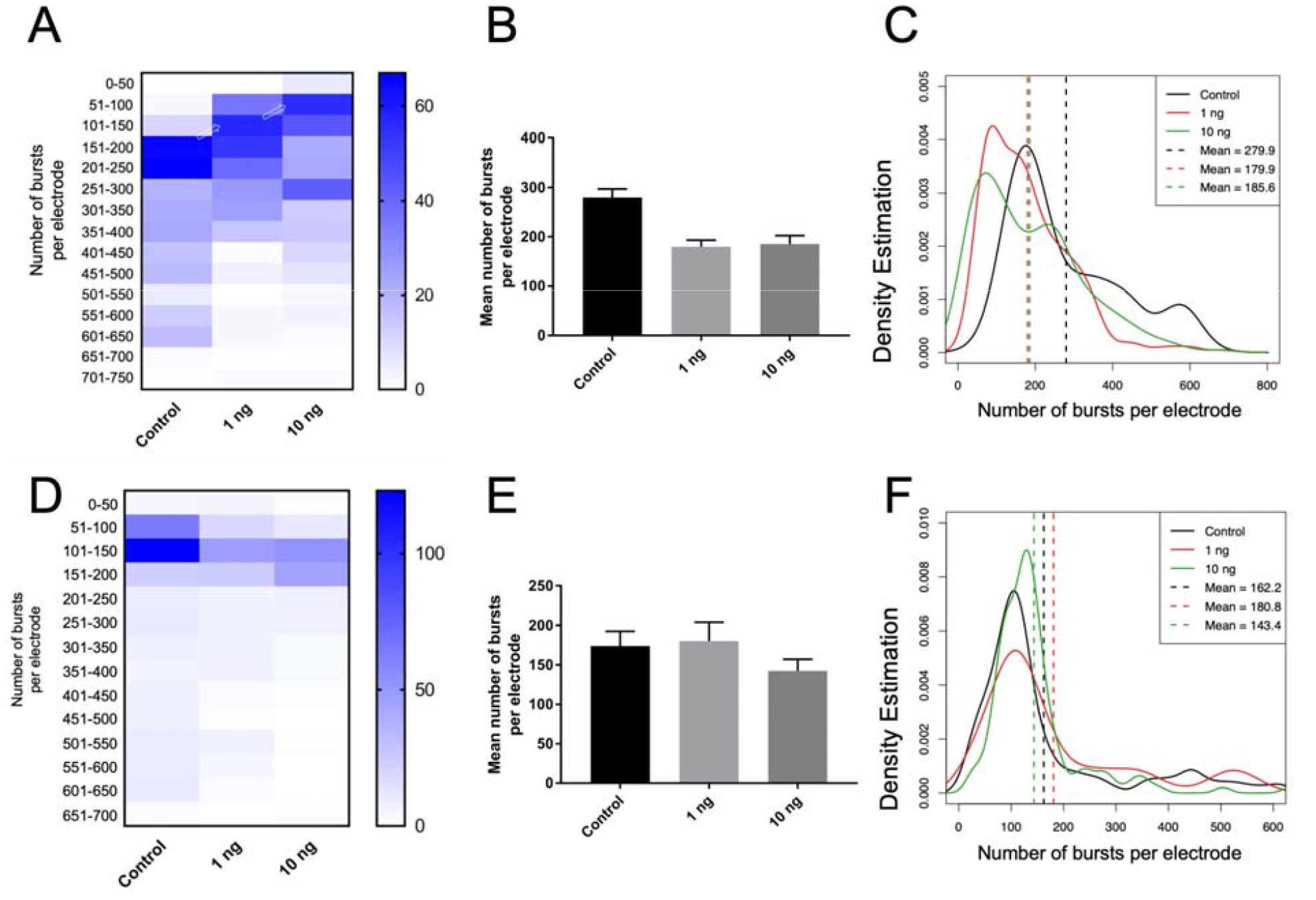
S1 reduced the number of bursts per electrode in neurons. 1 ng/mL and 10 ng/mL recombinant S1 (A-C) or S2 (D-F) were used to treat neurons on day 0. (A) The y-axis of the heatmap indicates the number of bursts per electrode, and the heat represents the number of electrodes (n). The arrows point to the shift in the number of bursts caused by 1 ng/mL or 10 ng/mL of S1. (B) The bar graph shows the mean number of bursts in each S1 treatment condition (p < 0.0001 for both 1 ng/mL and 10 ng/mL S1 compared to the control by unpaired t-test) (C) The density plot shows a shift in the distribution in wells treated with S1. The dotted line indicates the mean. (D) 1 ng/mL and 10 ng/mL S2 did not reduce the number of bursts per electrode. (E) 1 ng/mL S2 did not reduce the number of bursts compared to the control (p = 0.69), while 10 ng/mL S2 caused a borderline reduction in the number of bursts (p = 0.03).(F) The density plot shows that both concentrations of S2 did not cause a shift in the distribution compared to the control.

### Mature neurons were not affected by S1

Our next question was whether the S1 subunit affects mature neurons if cells are exposed to S1 later in the developmental course. The experiment was carried out with the same paradigm, except for the timeline of the S1 exposure. We treated the neurons with the same concentrations of S1 protein on day 12 and recorded the activities after the S1 exposure for 7 consecutive days. However, no noticeable reduction of burst activities was found between the S1-treated wells and the control (Figure 3). Therefore, the data suggest that the S1 subunit only affects neurons if cells are exposed early in the developmental stage.

**Figure 3.**
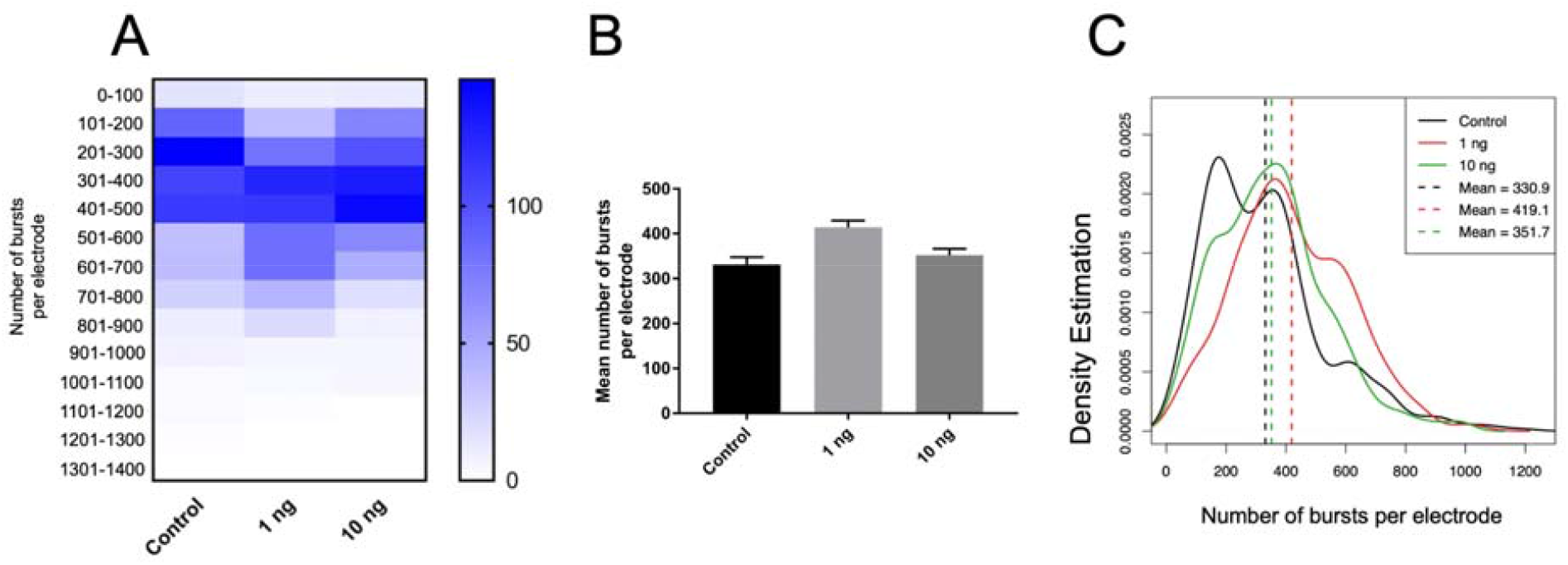
S1 did not reduce burst activities when the neurons were treated later in the developmental course. (A) Neurons were treated with 1 ng/mL S1 and 10 ng/mL respectively on day 12. The heatmap does not show a reduced burst number per electrode for both treatment groups compared to the control. (B) The bar graph shows that the 1 ng/mL S1 did not reduce the mean number of bursts per electrode compared to the control (p < 0.0001 by unpaired t-test). 10 ng/mL S1 also did not reduce the mean number of bursts per electrode (p = 0.07 by unpaired t-test). (C) The density distribution shows that both 1 ng/mL and 10 ng/mL did not cause a shift in distribution compared to the control.

### Human anti-S1 antibody rescued the neuronal phenotype caused by S1

To minimize the possibility that our previous observation was an artifact and the neuronal phenotype caused by S1 was in fact due to other uncharacterized factors, we conducted a rescue experiment to determine whether this neuronal phenotype can be reverted. A human monoclonal anti-S1 antibody (AS35 ACROBiosystems) was obtained and S1 was neutralized by the antibody before administering into neurons on day 0. The 1 ng/mL S1 that was neutralized by antibodies did not yield a significant reduction compared to those of the control (p = 0.77), while the regular S1 treatment on day 0 did decrease the burst activities (Figure 4). The data and p values strongly suggest that the anti-S1 antibody rescued the effect of S1 on bursting activities and the previous finding about S1’s ability to decrease neuron’s burst activities was unlikely to be an artifact. To conclude, the rescue experiment provides strong evidence that S1 was able to reduce burst activities if cells were exposed early in the developmental course.

**Figure 4.**
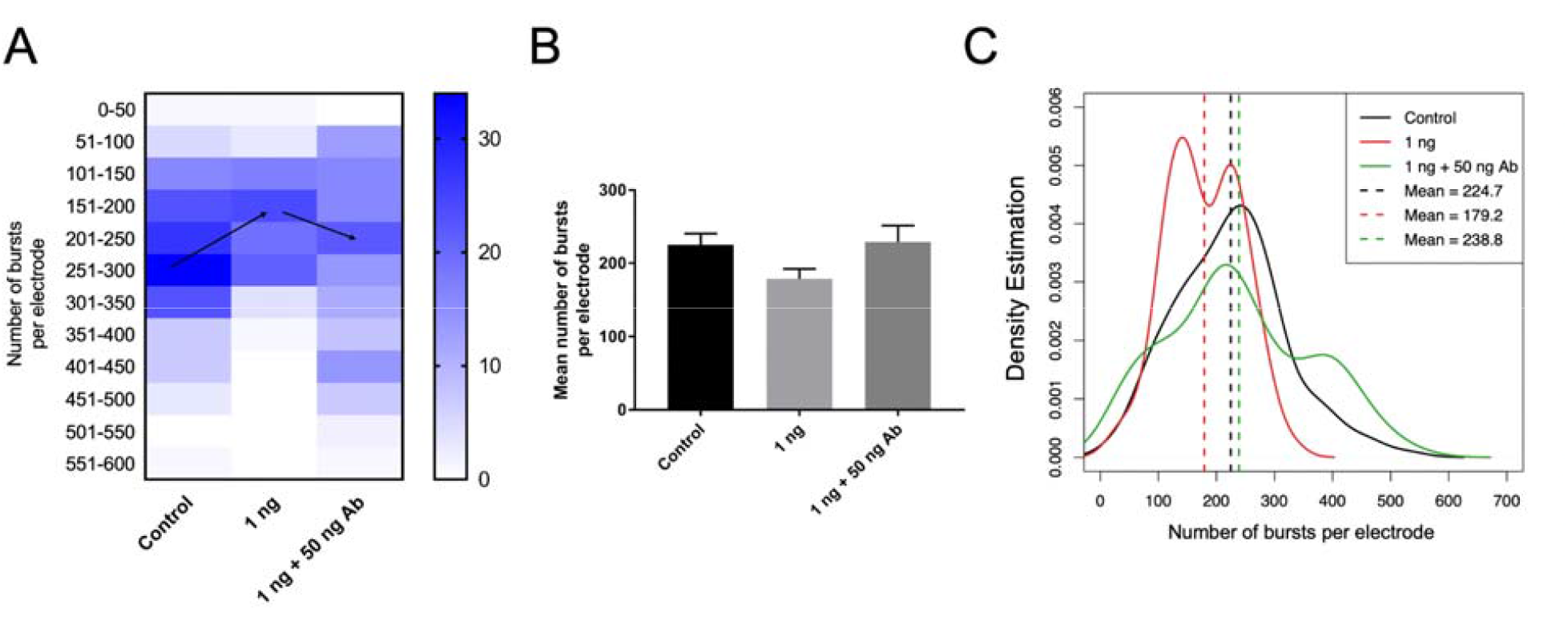
Human monoclonal anti-S1 antibody rescued the effect caused by S1. (A) 1 ng/mL S1 protein was neutralized by human anti-S1 antibody (50 ng/mL) at room temperature for 3 hours before administering into the culture media at day 0. (A) The heatmap confirmed that 1 ng/mL S1 reduced the number of bursts per electrode, and the antibody treatment reverted the effect of S1 to that similar to the control (indicated by the black arrow). (B) The bar graph shows that the 1 ng/mL S1 reduced the mean number of bursts per electrode (p < 0.0001 by unpaired t-test) and the neutralized S1 did not show a significant reduction (p = 0.77 by unpaired t-test). (C) 1 ng/mL S1 treatment (red) caused a left skew in the distribution while the S1 + antibody treatment (green) reverted the distribution to the curve similar to the control.

### The Receptor-Binding Domain (RBD) of S1 Is Responsible for Reducing Burst Activities

Despite the strong evidence that S1 reduced burst count during early treatment and that the effect was reversible by the human anti-S1 antibody, it is unclear whether the full-length or a specific domain of S1 is responsible for the burst reduction. Therefore, we hypothesized the receptor-binding domain (RBD) on S1 may be responsible for burst reduction, and we obtained purified recombinant RDB (Arg319 - Phe541) in order to test the hypothesis (Figure 1). The experimental design followed the same experimental paradigm in which neurons were treated with 1 ng/mL RBD on day 0. RBD concentrations were then maintained at the same concentration during the whole culture period. We identified a clear reduction in burst activities that are comparable with the S1 data (Figure 5). The result strongly suggests the fact that the RBD alone is sufficient to reduce burst activities.

**Figure 5.**
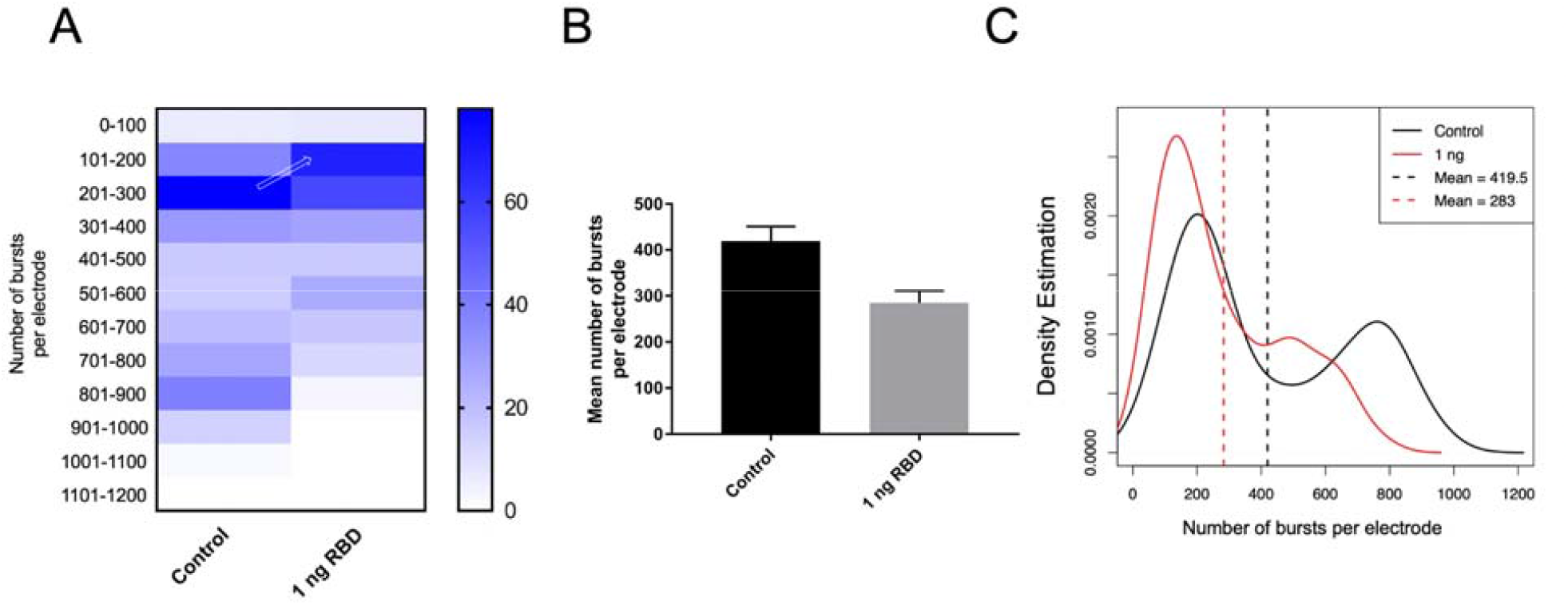
The Receptor Binding Domain (RBD) of S1 reduced the number of bursts per electrode in neurons. (A) 1 ng/mL purified RBD was used to treat neurons on day 0. The heatmap indicates a shift in the burst counts per electrode. (B) The bar graph shows a significant reduction in the mean number of bursts per electrode (p < 0.0001 by unpaired t-test). (C) RBD caused a shift in distribution (the dotted line represents the mean) in the density plot.

## Discussion

Our study shows a clear causal relation between the SARS-CoV-2 S1 protein and burst patterns in neuronal populations *in-vitro*, and this correlation can be rescued by antibody treatment. We further demonstrated that the receptor-binding domain might be responsible for the reduction of neuronal signals. Traditionally, the understanding of the pathogenicity induced by viral diseases is thought to be due to cell damages that are induced by the full viral replication cycle (involving attachment, entry, synthesis, and release). The full replication cycle destroys the host cells and releases more virions. In this process, tissue damage is induced which may recruit immune cells and induce inflammation. Here, we show a novel aspect of SARS-CoV-2 in a multi-well MEA model: Without the full infection by the SARS-CoV-2 virion, S1, alone, is sufficient to cause the phenotype of reduction in burst numbers. Furthermore, we show that the receptor-binding domain alone is sufficient to reduce burst activities, shedding new insights into the mechanism of SARS-CoV-2 that changes the cell’s phenotype beyond the traditionally understood mechanisms.

The mechanism of SARS-CoV-2, specifically the role of the S protein, and its influence on the neuronal system is still unclear and requires more investigation. Within the respiratory system, the COVID-19 infection is associated with a decreased total lung capacity, also resulting in overall respiratory failure and endothelial damage that disrupts pulmonary vasoregulation (Papa et al., 2021). Connecting to brainstem affection, evidence of acute respiratory failure as a result of COVID-19 from neurotropism of the brainstem by SARS-CoV-2 also leads to pulmonary damage, with the strain invading the brain through axonal transport via the olfactory nerves (Papa et al., 2021; Bauer et al., 2022). However, overall, there are limited findings about the neuronal impacts of SARS-CoV-2 as it travels within the lung-to-brain axis to alter neural development and respiratory function. To further understand the mechanisms and impacts of SARS-CoV-2 on the central nervous system (CNS), more research and tools are urgently required. Our study may provide the scientific community with a new instrument to study the viral impact on neurons, breaking it down to each viral protein part and assessing their influence on neurological phenotypes.

One of the more elusive aspects of COVID-19 infection is a subsequent phenomenon known as chronic post-COVID, more commonly known as long COVID (Sykes et al., 2021). Chronic post-COVID is described as prolonged symptoms after being infected by the COVID-19 virus, with symptoms lasting weeks, months, or even years after infection (Mahase., 2020). Chronic post-COVID is seen to be prevalent in patients who have been infected by the COVID-19 virus; Patients may still develop chronic post-COVID despite the seriousness of the infection or the degree of treatment received (Crook et al., 2021). There is no known mechanism for chronic post-COVID, only a theorized list of potential contributors such as post-COVID infection, organ damage, or chronic inflammation which may be caused by impaired ACE2 receptors, and traumatic effects of being in critical care (Verdecchia et al., 2020). Though further evaluation is needed, there seems to be an association between COVID recovery, specifically in a hospital setting, and deterioration of patient-reported quality of life. Many of the reported symptoms of chronic post-COVID are largely neurological and may require further evaluation or long-term care from medical professionals. There is no evidence that at a microbiological level, patients suffering from chronic post-COVID retain any trace of the virus within host cells, rather they only retain antibodies correlating to their stage of recovery (Proal and VanElzakker, 2021). Research seems to suggest that there is a correlation between the severity of the illness present and the severity of chronic post-COVID symptoms, meaning that those who were either asymptomatic or not critically infected with the virus have a better chance of not developing chronic post-COVID while those who were hospitalized are nearly twice as likely to report symptoms of chronic post-COVID (Raveendran et al., 2021). Oftentimes, patients with long-term respiratory issues also complain of chronic fatigue and headaches while those with gastrointestinal issues are more likely to present elevated temperatures. Our research suggests the likelihood that the spike protein of SARS-CoV-2 has the potential to induce neurological impacts that were not previously understood, which may partially explain some aspects of the chronic post-COVID symptoms associated with the central nervous system. However, further investigation is required to fully establish this direct causal relationship in patients.

The SARS-CoV-2 RBD is a common target for antibodies and vaccines (Min and Sun, 2021). SARS-CoV-2 RBD is made up of two main structures, the core and receptor-binding motif (RBM). The RBM is the main fragment of the RBD that contacts the ACE2 receptor (Freund et al., 2015; Li et al., 2005). The RBM of the RBD contacts the underside of a small part of the ACE2 receptor to facilitate cell entry (Lan et al., 2020; Shang et al., 2020). Here, we offered functional insight into the RBD domain of S1 and its potential importance in affecting neuronal activities. However, more investigation is required to further understand the RBD and the mechanism of how RBD alone is sufficient to affect neuronal activities.

Our study shows that by neutralizing S1, the neuronal burst activities recover to the level of the control. The human monoclonal anti-S1 antibody functions by binding to the RBD on the spike protein and preventing it from binding to ACE2 (Barnes et al., 2020). Our antibody rescue experiment not only confirms the role of S1 in reducing burst activities, but also emphasizes the protective role of anti-S1 antibodies and the importance of RBD in affecting neuronal phenotypes. This finding is consistent with the clinical observations (Hurt et al., 2021; Min and Sun., 2021; Ho et al., 2020). Additionally, it may also be interesting to ask the question of whether antivirals against SARS-CoV-2 also have similar protective mechanisms against S1’s impact on neurons. Overall, our study offers new insight into the understanding of the mechanism of SARS-CoV-2 spike proteins beyond their traditionally known roles of viral attachment and entry. This finding may inform important aspects about the biology of SARS-CoV-2, patient care strategies, or even future vaccine or antiviral designs.

